# Recombinant Group A Carbohydrate backbone embedded into Outer Membrane Vesicles is a potent vaccine candidate targeting Group A Streptococcus from *Streptococcus pyogenes* and *Streptococcus dysgalactiae* subsp. *equisimilis*

**DOI:** 10.1101/2021.11.12.468441

**Authors:** Sowmya Ajay Castro, Sarah Thomson, Azul Zorzoli, Benjamin H Meyer, Mark Reglinski, Helge C. Dorfmueller

## Abstract

**Background:** Group A Streptococcus (GAS) are responsible for a wide range of human-exclusive infections, annually killing more than 500,000 people. Antibiotic resistance incidence of invasive GAS tripled in the past decade and emphasises the need to develop a universal GAS vaccine. We have produced, for the first time, a recombinant polyrhamnose backbone (pRha), a validated universal GAS vaccine candidate. *E. coli* outer membrane vesicles (OMVs) carrying pRha were investigated for their immunogenicity and efficacy in an animal model.

**Methods:** OMVs decorated with pRha were administered to C57BL/6J mouse and rabbit models. Flow cytometry, ELISA, Luminex, immunofluorescence microscopy and serum bactericidal assay assays were conducted to investigate the ability of pRha-specific antibodies to recognise and kill clinical (hypervirulent) GAS strains.

**Results:** Our results suggest that pRha-OMVs induce specific antibodies which recognise Group A Carbohydrate (GAC) from *S pyogenes* and *S. dysgalactiae* subsp. *equisimilis*. Increased IgG levels correlate with increased bactericidal killing of the hypervirulent GAS M89 strain. Elevated IL-17a from pRha-OMV-immunised splenocytes indicates possible stimulation of long-term memory immune cells.

**Conclusion:** We are the first to report efficacy and potency of this unique, exogenously produced polysaccharide, pRha, in the induction of humoral-mediated immune responses to GAS.

**Topic:** *Streptococcus pyogenes*, immunoglobulins, polysaccharides, opsonophagocytosis, acute rheumatic fever, M protein, invasive Group A Streptococcus, hyaluronic acid

## Introduction

*Streptococcus pyogenes*, frequently described as Group A Streptococcus (GAS), is a beta-haemolytic Gram-positive human exclusive bacterium causing a wide range of illnesses in both healthy and immunocompromised individuals. Breaching of immune barriers by GAS leads to acute suppurative effects in the skin and pharyngeal epithelia, causing localised skin infections, tonsillitis and enlarged lymph nodes in the throat [1]. Prolonged prevalence of GAS in the epithelial cells has proven to lead to life-threatening complications such as streptococcal toxic shock syndrome, necrotising fasciitis and, importantly, post-streptococcal autoimmune sequelae [2]. One of these major autoimmune sequelae is rheumatic heart disease, which kills >300,000 patients worldwide [1, 3]. Incidence of antibiotic resistance has tripled in the past decade, and the common penicillin-based antibiotics only eradicate ∼65% of all tonsilitis cases caused by GAS infections [4, 5].

Identification of GAS is based on the detection of its main surface antigen, the Group A Carbohydrate [GAC]. This consists of a polyrhamnose (pRha) backbone (→3)α-Rha(1→2)α-Rha(1→) with an immunodominant N-acetylglucosamine (GlcNAc) side chain present on every α-1,2-linked rhamnose [6-8]. Recent studies have reported the presence of negatively charged glycerol-phosphate onto approximately every fourth GlcNAc sidechain [9]. Earlier research proposed GAC as a potential antigen due to i) its abundant presence in the cell wall contributing ∼50% of GAS by weight, ii) its conservation across all >200 GAS serotypes and sequenced GAS genomes [10], and iii) the absence of cross-reactivity between antibodies raised against endogenously-produced pRha and human cardiac antigen [11, 12].

GAC antibodies were frequently found at high-titres in serum samples, directly correlating with the presence of GAS in the throat of children [13], and anti-GAC antibody titres persist in patients with rheumatic valvular disease [14]. Moreover, a strong body of evidence shows that injecting purified or synthetic components of GAC conjugated to a carrier protein elicits an immune response that confers protection against multiple GAS serotypes [13, 15]. However, the anti-GAC antibodies raised from immunisation have safety concerns: N-GlcNAc-specific antibodies against the sidechain cross-react with human tissues [16, 17]. Importantly, this was not seen in antibodies raised against the pRha backbone alone [11, 12] suggesting that pRha could be a safe and effective vaccine candidate.

Outer membrane vesicle (OMV)-based vaccine candidates produced in *E. coli* have been in development for more than ten years and have shown efficacy against several pathogens [18, 19]. Here, we show that recombinantly produced pRha-OMVs carry the correct pRha structure, as revealed by linkage analysis, identical to the natively-produced pRha in a GAS mutant strain that is deficient in the GlcNAc side chain [11, 12]. We show that *E. coli* pRha-OMV are immunogenic in a mouse model; moreover, the stimulated IgG antibodies are able to opsonise GAS isolates and *S. dysgalactiae* subsp. *equisimilis* (SDSE) and promote the killing of hypervirulent GAS strains. Increased IL-17a indicated the possible stimulation of cellular-mediated immune response in pRha-OMV vaccinated animals. We thus present an alternative route to recombinantly produce a universal GAS OMV-based vaccine candidate, removing the requirement to isolate the pRha chemically or enzymatically from GAS cells.

## Materials and Methods

### Bacterial strains and growth conditions

A single *E. coli* colony was cultured in 10 mL LB medium supplemented with 150 µg/mL erythromycin, grown at 37°C, 200 rpm overnight and used to inoculate 1 L medium, grown for 18 h. GAS isolates were grown at 37 °C, 5% CO_2_ in THY. *E. coli* and GAS isolates were used as overnight cultures for Western blot analysis or used at 1:100 to O.D_600_ 0.4 for serum bactericidal assay, flow cytometry and ELISA assays.

### Ethical approval

Mice studies were approved by the University of Dundee welfare and ethical use of animals committee and conducted in accordance with the UK Home Office approved project licence [PPL PEA2606D2]. The experiments involving rabbits were conducted in association with Davids Biotechnologie GmbH, Germany.

### Purification of pRha-OMVs

*E. coli* OMVs were isolated from CS2775 cells transformed with plasmids pHD0136 and pHD0139 [9]. Cells were harvested by centrifugation (4,500 RPM, 30 min, 4°C). The supernatant was filtered (0.45μm) and centrifuged (Type 45Ti, 45,000 RPM, 4 hrs, 4°C). The pellets were weighed, resuspended in PBS, and protein content determined using the Bradford method. The OMVs were analysed for their average size distribution in the Zetaview nanoparticle tracking analysis instrument, and LPS levels assessed using the Limulus Amoebocyte Lysate (LAL) assay (Thermo Fisher, UK).

### Immunisation

Mouse Model: Five- to six-week-old female C57BL/6J mice were acquired from Charles River Laboratories, UK, and were acclimatised for 10 days prior to immunisation. Mice (n=6) were immunised subcutaneously with pRha-OMV (10 µg diluted in 100 μl of endotoxin free PBS/mouse/injection) or *E. coli* OMV (n=6) on day 0, immediately following tail bleed for pre-immune sera. Identical booster injections were administered on day 21 and day 49. Animals were euthanised by CO_2_ asphyxiation and bled by cardiac puncture on either day 70 or day 84. For adjuvant studies, Imject Alum (Sigma) was used with 10 µg of pRha-OMV (1:1 ratio); adjuvant with PBS only (1:1 ratio) served as a negative control. PBS/Alum vaccinated mice (n=3) were used for baseline measurements.

Rabbit Model: For New Zealand White rabbit immunisation, the animal (n=1) was bled for pre-immune serum prior to immunising with the purified OMV pRha (150 µg diluted in 150 μl of endotoxin-free PBS/injection) on day 0. Booster injections were administered on days 14, 28, 42 and 56, at 100 µg in 100 μl of endotoxin-free PBS/injection. The animal was culled by CO_2_ asphyxiation and bled by cardiac puncture. Final bleed serum was taken on day 63 and affinity-purified (0.53 mg/ml) by Davids Biotechnologie GmbH.

### Flow cytometry

Antibody binding to GAS and *E. coli* cells was adapted from [20]. Bacterial cells (2 × 10^6^ CFU) were resuspended in nonspecific human IgG (Sigma, UK) for one hour on ice, then washed with PBS and diluted 1:100 with immunised mice serum (PBS or pRha-OMV) overnight at 4°C. Cells were washed twice with PBS and stained with 1:250 dilution of Alexa fluor 488-conjugated goat anti-mouse IgG (Thermo Fisher, UK), incubated for 20 minutes at 4°C in the dark. Cells were washed twice (PBS) and fixed with 500 μl of 4% PFA and analysed by flow cytometry (BD Bioscience). A total of 10,000 events were acquired and data were analysed using FlowJo software version 10.6.2.

### ELISA

For antigen-antibody interaction, plates were coated with 50 μl/well of 1 μg/ml pRha-OMV/negative control OMVs or *S. pyogenes* cells adapted from [21, 22]. Briefly, bacterial cells at OD_600_ of 0.4, from the overnight culture, were resuspended (1:100) in PBS. Mouse serum from immunised animals was added, 50 µl/well at 1:1,000. Bound antibodies were probed using 50 μl/well of 1:1,000 of anti-mouse IgG HRP (Sigma–Aldrich) and 75 μl/well of tetramethylbenzidine substrate (Sigma–Aldrich). The reaction was stopped using 75 μl/well of 1 M H_2_SO_4,_ and absorbance read at 450 nm.

### Luminex

Immunoglobulins in mouse sera were quantified using a Milliplex Multiplex assay (Merck Millipore, UK). Immunoglobulin beads (IgA, IgG1, IgG2a, IgG2b) from Mouse Antibody Isotyping 7-Plex ProcartaPlex were used as analytes and the samples were prepared according to manufacturer’s instructions (Thermo Fisher, UK).

### Immunoblot analyses

Antisera from pRha-OMV vaccinated animals were tested for binding to *E. coli* cells or GAS by Western blot analysis using 12.5% acrylamide PAGE gels (Thermo Fisher, UK). *E. coli* cells and OMVs were used from overnight cultures. Overnight-grown GAS cells were washed with PBS and incubated with 6 μl of PlyC (0.7 mg/ml) for one hour (37°C, 300 RPM) and centrifuged for 14,000 RPM, 5 mins. The resulting pellets were resuspended in 2x SDS-PAGE loading dye and subjected to Western blot analysis (Invitrogen, UK). The PVDF membranes were blocked with 5% non-fat dried milk in Tris-Buffered Saline, 0.1% Tween® 20. Immunised mouse sera were used at 1:1000 dilution to probe the blots (overnight, 4°C, 20 RPM) followed by goat anti-mouse IgG HRP at 1:1000 dilution for two hours at 4°C, 20 RPM. Rabbit anti-GAC antibodies (Abcam, UK) with goat anti-rabbit IgG HRP (Abcam, UK) (1:2,500) were used as a positive control. Antibody binding was visualised using Clarity Western ECL Substrate (Biorad, UK) and viewed under the Gel Doc imaging system (GeneSys software, Syngene).

### Isolation and *ex vivo* re-stimulation of splenocytes

Isolation of mouse spleens and re-stimulation with antigens was conducted according to published procedures [23]. Briefly, spleens were rinsed and flushed, using 21G needles, with 5 ml of RPMI 1640 media (supplemented with 100 U/ml Pen-Strep (Sigma, UK), 200 mM L-glutamine (Gibco, UK) and 10% heat-inactivated foetal bovine serum (Sigma, UK)). The dispersed cells were centrifuged (300xg, 10 min) and red blood cells were lysed in lysis buffer (Sigma, UK). Cells were washed and centrifuged again by the addition of 10 ml of PBS (300xg, 10 min). Pellets were resuspended in complete RPMI 1640 media and adjusted to 7.5×10^6^ cells/ml; 1.5×10^6^ cells [200 µl] were added to each well of a 96 well plate followed by 50 µl of 1 µg/ml pRha-OMV and incubated for three days. Cells were pelleted (400xg, 10 min) and supernatants subjected to ELISA analysis for IL-17a (Thermo Fisher, UK).

### Microscopic analysis

GAS M89 cells were grown as mentioned above. Briefly, overnight culture of M89 strains were washed twice with PBS (10,000 RPM, 5 minutes). Washed cells were stained overnight at 4°C with pre-rabbit or pRha-OMV post immune sera at 1:100 dilution. Prior to adding secondary antibody (goat anti-rabbit FITC in 1:50) the cells were washed twice with PBS. The FITC-stained cells were mounted and viewed using a DeltaVision microscope; images were deconvoluted and analysed using softWoRx imaging system.

### Serum Bactericidal assay (SBA)

SBA assays were carried out according to [24, 25]. Briefly, in a 96-wellplate 25 μl of HBSS medium containing CaCl_2_ and MgCl_2_ (Gibco), followed by 25 μl of rabbit antiserum (1:100 dilution). The antibodies were incubated with 12.5 μl of M89 (2×10^5^) cells in the presence of activated or inactivated baby rabbit complement (37°C, 3h, 5%(v/v) CO_2_), then 12.5 μl of mixed cells were spotted at the start of the incubation (T0) and at the end of incubation (T3) onto sheep blood agar plates (Oxoid, UK) and incubated overnight (37°C, 5%(v/v) CO_2_). Colonies were counted from duplicate wells and analysed for the percentage of bacterial survival.

### Statistical Analysis

Data were analysed using GraphPad Prism Version 8 (La Jolla, CA, USA). One-way analysis of variance [ANOVA] with Bonferroni’s correction, two-tailed T-test followed by Dunn’s post-test were used to test the statistical significance. FlowJo analysis software (FlowJo, USA) was used to plot the geometric mean fluorescence intensity (gMFI) of flow cytometric data.

## Results

### Recombinantly produced polyrhamnose-OMVs carry the antigen on Lipid A

The polyrhamnose (pRha) carbohydrate antigen with the immunospecific repeat unit of two rhamnose sugars, linked via the alpha1,2-1,3 repeating unit, is found in GAS, GCS and *S. mutans*, and also in newly emerging isolates from *S. dysgalactiae* subsp. *equisimilis* [11, 26-28]. We recombinantly expressed the gene cluster from *S. mutans sccA-G* and *S. pyogenes* in *E. coli* [9, 29]. Our analysis revealed that the pRha thus produced is deposited on the outer membrane Lipid A, transferred onto it in the inner membrane via the O-antigen ligase WaaL [Fig.1A & Supplementary Fig.1A] [30, 31]. We isolated OMVs that were either decorated with the pRha (pRha-OMV) [Fig.1B] or lacking the pRha (empty OMVs) and analysed them using Group A Carbohydrate specific antibodies. Only the pRha-OMVs were detected by the antibodies, confirming the decoration of the OMVs with pRha [Fig.1B]. The purified OMV’s were quantified and tested for LPS content using the LAL kit, confirming the endotoxin levels at 0.7 EU/ml [Supplementary Fig.1B]. The OMV average particle sizes were in agreement with previously reported *E. coli* OMVs, with a diameter range of 50–250 nm [Supplementary Fig.1C] [32].

**Figure 1:**
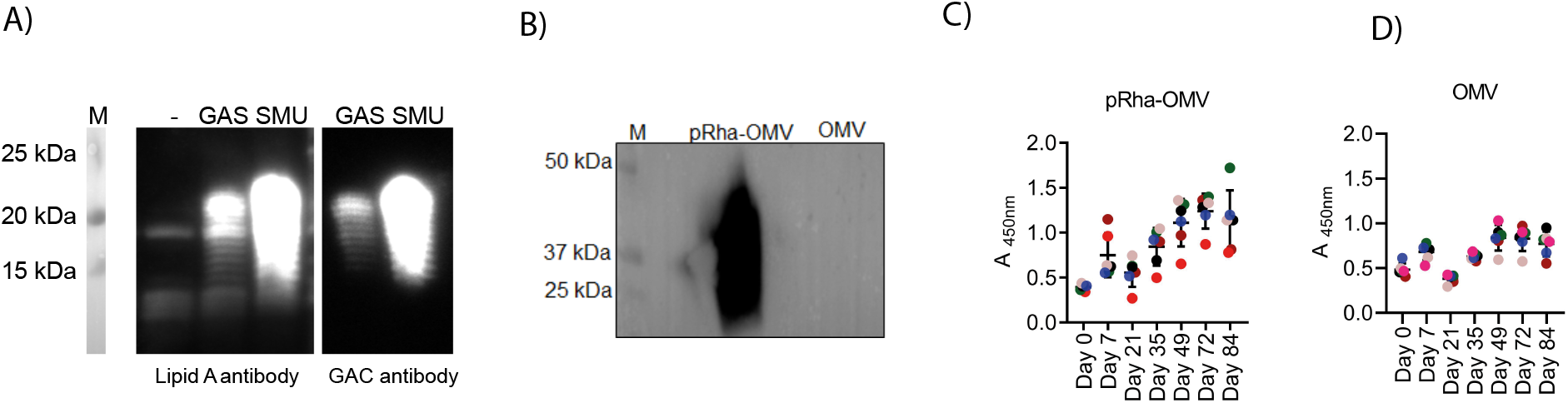
Purification and Immunisation of recombinantly produced pRha-OMV: A) Western blot of pRha produced in *E. coli* cells from the GAS and *Streptococcus* mutans (SMU) gene clusters and probed with Lipid A antibody and GAC specific antibody. B) Representation of immunoblot analysis of recombinantly produced *E. coli* producing pRha-OMV and *E. coli* OMV were stained for 1:1000 Rabbit anti-GAC antibody followed by goat anti-IgG HRP. Molecular mass markers are given in kilodaltons. C) pRha-OMV specific IgG antibodies were measured by coating the 96-well plate with MEGA cells (*E. coli* cells expressing longer chain of GAS pRha) and analysed using sandwich ELISA. The anti-pRha-OMV sera (C) or OMV vaccinated mice sera (D) collected at each stage of immunisation were tested against MEGA. Bound antibodies were detected using pre-titrated, biotin-conjugated antibody. Data shown are mean ± S.E.M. from technical replicates.

OMVs have been shown to induce IgG antibodies in animals [33] as well as in humans [34]. We immunised mice with either pRha-OMV or control OMVs and measured the levels of pRha-OMV-specific IgG by exposing the antibodies to *E. coli* control strain that produces a higher level of polyrhamnose [Fig.1C, D]. The pRha-OMV-specific IgG levels increased in pRha-OMV immunised animals compared to those in control animals (immunised with empty OMV), demonstrating increased binding of the anti-pRha-OMV antibodies to the polyrhamnose compared to anti-OMV titres [Fig.1C, D].

### pRha-OMV stimulated antibodies selectively bind to the pRha epitope

Final bleed sera from each group were pooled and analysed by flow cytometry for their ability to recognise the pRha antigen expressed on *E. coli* cells compared with control cells lacking pRha. Histograms for antibody deposition and extraction of gMFI from the flow cytometry data generated using pRha-OMV immunised animals clearly revealed increased IgG antibody deposition onto *E. coli* pRha-bound OMVs, compared to control anti-OMV IgG [Fig.2A,B], indicating that selective paratopes are expressed in the sera of the pRha-OMV vaccinated animals. Importantly, the gMFI for pRha-OMV was significantly higher than that of the negative control (OMV) on exposure to pRha positive *E. coli* cells suggesting strong immunogenicity of the pRha [Fig.2B]. This agrees with the observation that OMVs alone stimulated production of antibodies that recognise *E. coli* cells without pRha, and, to a lesser degree, *E. coli* decorated with pRha [Fig.2B]. Interestingly, the levels of cross-reactivity were similar for both species (∼50 gMFI units), supporting our interpretation that pRha-OMVs stimulate production mainly of antibodies against the pRha but also, to a lower extent, against the OMVs.

**Figure 2:**
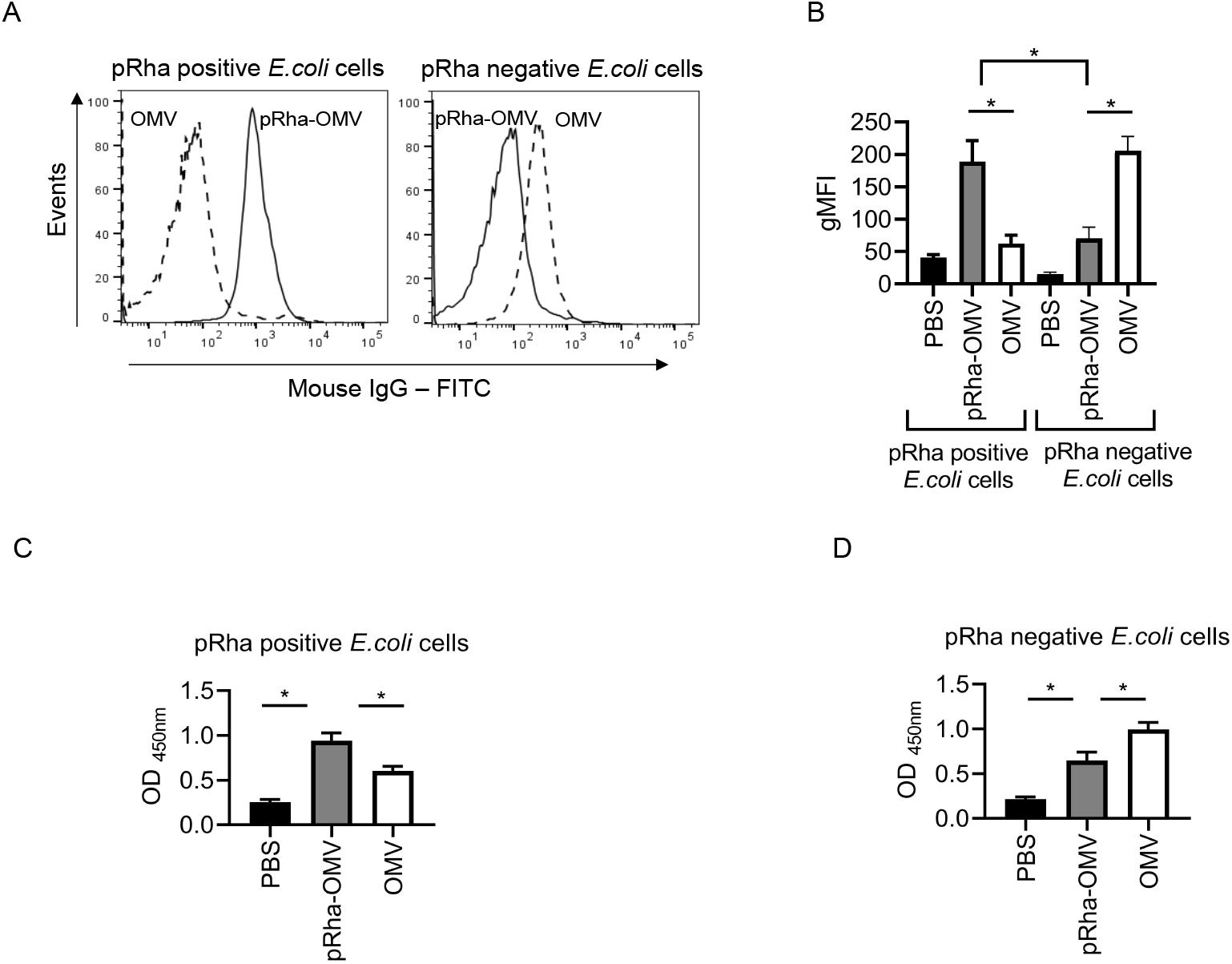
pRha-OMV induced IgG antibodies selectively binds to pRha positive *E. coli* cells: A) Representative histograms for antibody deposition on pRha positive *E. coli* cells or pRha negative *E. coli* cells stained either using pooled 1:100 antiserum from pRha-OMV immunised sera (open black line) or 1:100 anti-serum from OMV alone immunised sera (dashed line). The stained cells were probed with anti-mouse IgG FITC and read in flow cytometer analysis. (B) IgG deposition of pRha-OMV immunised or OMV alone immunised or PBS animal sera analysed using geometric mean fluorescence intensity (gFMI) by treating either with the pRha positive *E. coli* cells or pRha negative *E. coli* cells. C) Plates were coated with pRha positive *E. coli* cells or pRha negative *E. coli* cells (D) and probed with a 1:1000 dilution of murine sera recovered from pRha-OMV immunised or OMV immunised or PBS mice. Bound antibodies were detected using a 1:1000 dilution of HRP-conjugated goat anti-mouse IgG. Flow data shown are collected from at least 10000 events. Statistical analyses were conducted using ANOVA followed by Bonferroni post hoc-test *P<0.05. Data shown are mean±S.E.M. of three independent experiments.

Using ELISA, serum from the pRha-OMV-vaccinated animal showed significantly higher binding affinity for the pRha-bound *E. coli* than for OMV or PBS immunised animals [Fig.2C]. These data indicate that the pRha antigen on the *E. coli* cell was recognised robustly by the pRha-OMV-immunised animals. Control, OMV-immunised, animal sera showed background staining possibly as a result of anti-OMV activity in these animals. Although, the antibody response from the OMV vaccinated animals was lower towards the pRha producing *E. coli* cells [Fig.2C], the anti-OMV titre in these animals was significantly higher in negative control cells alone compared to anti-pRha-OMV titre groups [Fig.2D]. Immunoblot analysis of pRha-OMV immunised serum showed a strong signal in *E. coli* pRha-OMV lysate confirming the presence of GAC [Supplementary Fig 1D].

### Validation of IgG subtypes in pRha-OMV immunised mice

Previous research has shown a correlation between frequent exposure to polysaccharides and increased IgG subtypes in humans and animals [35-37]. Hence, we examined the effect of pRha-OMV on the IgG subtypes in mice (IgG1, IgG2a, IgG2b and IgG3) using Luminex analysis [Fig.3]. The Ig subtypes were analysed in pooled sera of the PBS and pRha-OMV immunised animals on day 49 (after the administration of two boosters) and on day 70 (after three boosters) [Fig.3]. Our results show a pattern of IgG subtype responses, revealing increased levels of total IgG2b [Fig.3C], IgG3 [Fig.3D] and IgG2a [Fig.3B], with lower levels of IgG1 [Fig.3A], from both PBS and anti-pRha OMV sera. However, the anti-pRha OMV IgG subtypes were significantly higher compared to PBS immunoglobulin levels and sustained until the end of the immunisation schedule (day 70).

**Figure 3:**
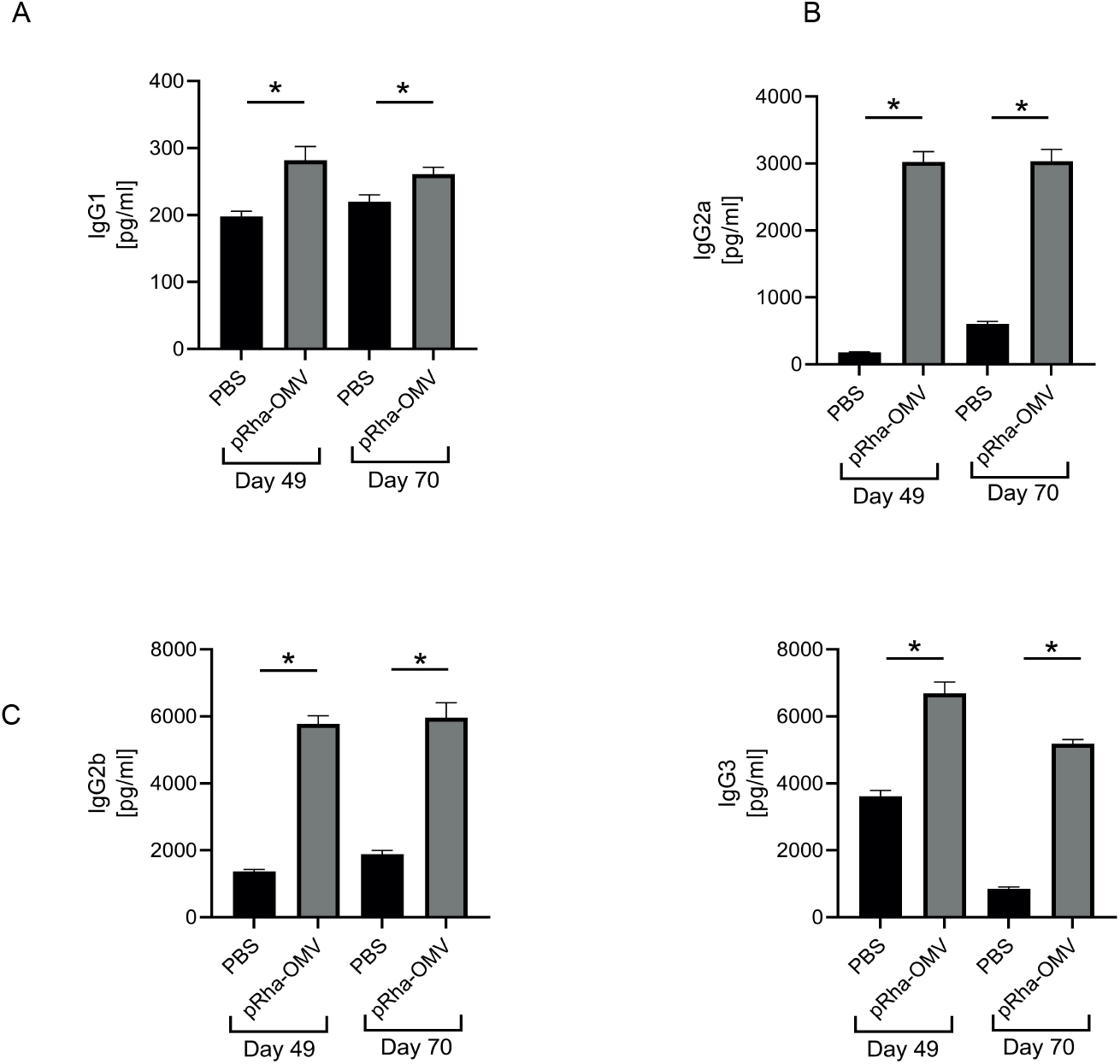
Quantification of Immunoglobulin subtypes against pRha-OMV in mice sera using Luminex analysis: A) IgG1 B) IgG2a C) IgG2b and D) IgG3 immunoglobulins were compared between day 49 (after two boosters) and day 70 (after three boosters) on the pooled pRha-OMV immunised mice sera or with PBS sera and analysed using Luminex. *p < 0.05 unpaired two-tailed T test (vs PBS). Data shown are mean±S.E.M. from technical replicates.

### pRha-OMV antibodies recognise clinical GAS isolates

The efficacy of the pRha-OMV antibodies was further investigated by testing their ability to recognise GAS isolates using flow cytometry and ELISA analyses. To assess the opsonisation of a wide range of clinical GAS serotypes, flow cytometry was performed using antibody titres from all groups. The gMFI obtained from the flow cytometry data showed significantly increased antibody deposition in sera of animals immunised with pRha-OMV compared to that in control immunised animals [Fig.4A]. However, not all GAS serotypes were recognised by the pRha-OMV IgG. Addition of aluminium adjuvant to the pRha-OMVs increased antibody deposition in mouse sera by recognising most of the GAS serotypes, except for M28 and M1_UK_ [Fig.4B]. The degree of IgG binding to the GAS isolates varied highly across all tested serotypes; the pRha-OMV with aluminium performed better than pRha-OMV serum alone [Fig.4A,B]. ELISA assays confirmed a similar response: pRha-OMV sera recognising GAS serotypes, including the dominant clade M1T1 (M1_UK_), a new *emm*1 *S. pyogenes* lineage [Fig.4C] [38].

**Figure 4:**
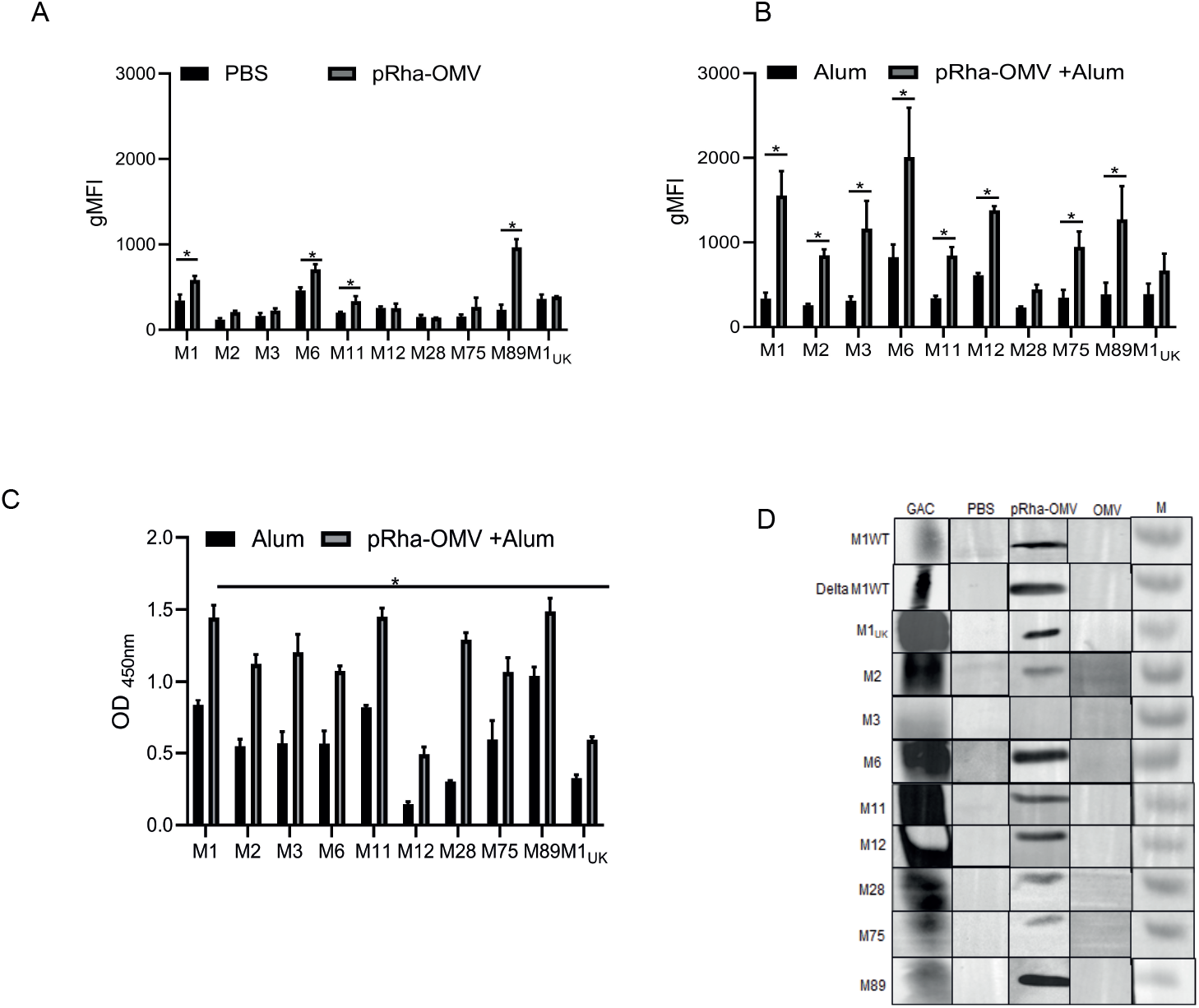
pRha-OMV vaccinated antibody deposition on GAS clinical isolates: A) Graph showing geometric mean fluorescence intensity (gMFI) extracted from flow cytometry analysis of clinical GAS isolates stained with either PBS or pRha-OMV pooled mice sera (1:100) followed by anti-mouse IgG alexafluro 488 channel B) Graph showing gMFI extracted from flow cytometry analysis of clinical GAS isolates stained with either alum or pRha-OMV with alum pooled mice sera (1:100) followed by anti-mouse IgG alexafluro 488 channel C) Whole cell ELISA analysis conducted to measure the antiserum from pRha-OMV with alum (1:1000) or antiserum from alum (1:1000) binding to clinical GAS strains D) Immunoblot analysis of clinical GAS isolates stained using pooled mice sera from the animals immunised with either PBS or pRha-OMV or OMV were used to probe the GAS lysates at 1:1000 dilution. The GAS lysates were further probed with anti-mouse IgG HRP (1:1000). Anti-Group A Carbohydrate antibody (GAC) was used as a positive control (1:1000). Statistical analyses were conducted using ANOVA followed by Bonferroni post hoc-test *P<0.05 (PBS vs pRha-OMV or Alum vs pRha-OMV+Alum). Data shown are mean±S.E.M. of three independent

GAS isolates were probed with a commercially available Group A Carbohydrate antibody and compared to pRha-OMV and negative control (OMV alone or PBS-immunised) mouse sera. GAS isolates show an intense band at ∼25 kDa on Western blots probed with sera from pRha-OMV mice, confirming the specificity of the pRha antibodies. The band was absent in the control OMV sera. The positive control (GAC antibody) displayed a stronger signal representing the presence of complete GAC components (including GlcNAc/glycerol-phosphate sidechain) [Fig.4D].

### pRha-OMV sera stimulate IL-17a production in murine splenocytes

We investigated the effect of the pRha-OMV immunogen on IL-17a production in splenocytes. IL-17a, a key cytokine mediator, contributes to activation of T cells by mediating protective innate immunity against GAS [39]. Increased IL-17a levels were detected in pRha-OMV vaccinated splenocytes restimulated with pRha-OMV antigen, compared with splenocytes from the PBS group [Fig.5A]. The data suggest that pRha-OMV induces a humoral mediated immune response, and also a cytokine-mediated cellular immune response [39, 40], potentially via carbohydrate-specific T helper cells [41].

**Figure 5:**
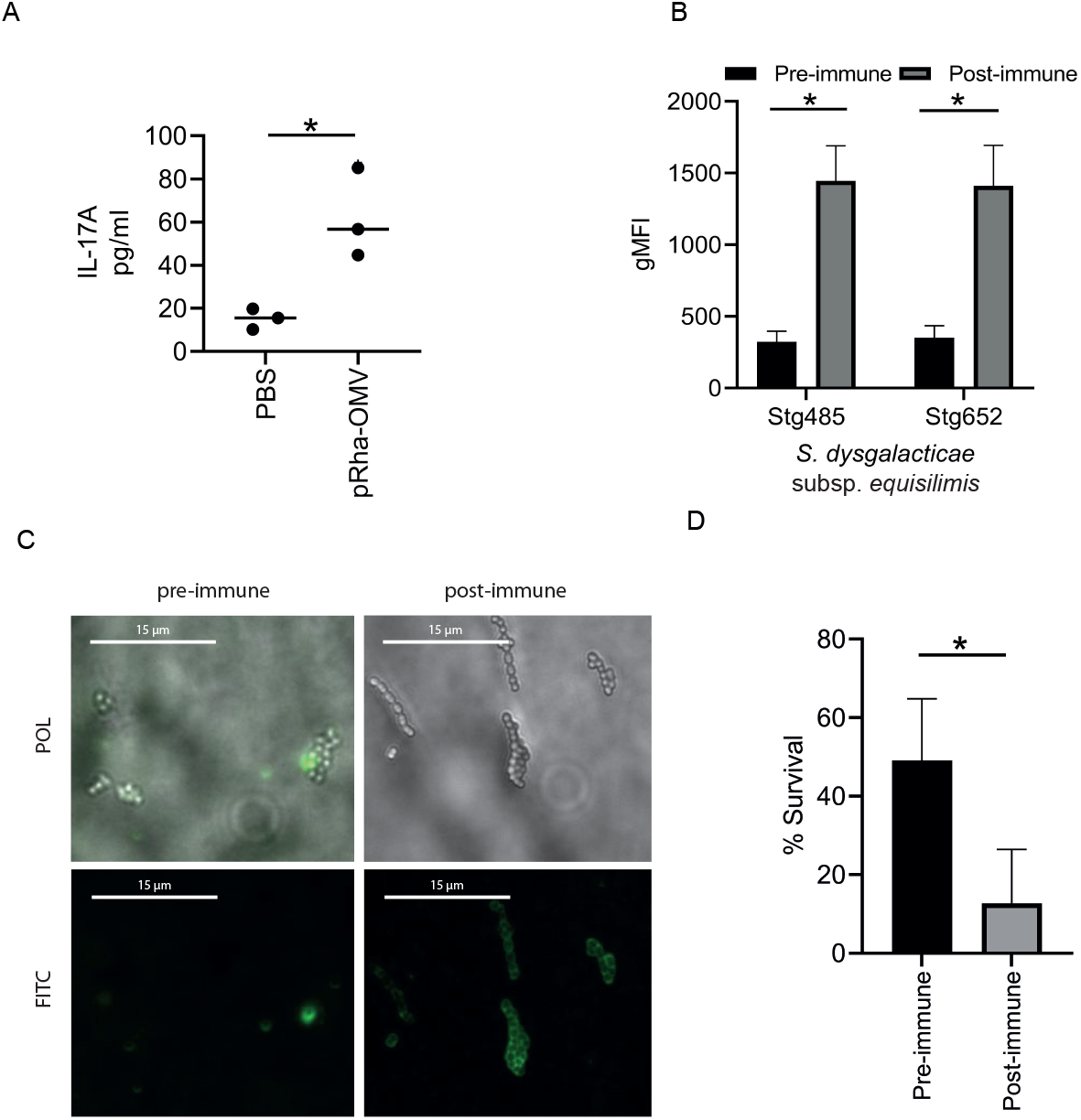
Rabbit immunised pRha-OMV IgG promotes killing of the hypervirulent strain M89: A) Graph showing the cytokine mediator IL-17a measured in the supernatants of individual mice splenocytes (PBS or pRha-OMV) restimulated with recombinant OMV pRha antigen (10 µg/ml). Results displayed as mean ± SEM from technical replicates B) Antibody deposition measured using a flow cytometry assay on *Streptococcus dysgalacticae* subsp. *equlisimils* containing GAC and lacking Group G Carbohydrate (GGC) (Stg485 and Stg652) stained in 1:1000 rabbit antiserum from pre-rabbit sera and pRha-OMV immunised sera. C) Representative images of immunofluorescent staining of hypervirulent strains M89 stained with either with pre-immune rabbit sera or pRha-OMV immunised rabbit sera (1:1000) followed by goat anti-Rabbit IgG FITC (1:50) channel. Polarised (POL) and FITC channel were used to document the images using deltavision microscopy D) Percentage of survival of M89 hypervirulent GAS strains were analysed using antiserum from the pRha-OMV immunised or pre-immune rabbit in the presence of 5% baby rabbit serum. *p < 0.05 unpaired two-tailed T test (PBS vs pRha-OMV or pre-immune sera vs post-immune sera). Data shown are mean±S.E.M. of three independent experiments for B, D.

### pRha-OMV immunised rabbit serum recognises GAS from *S. dysgalactiae* subsp. *equisimilis* and kills the hypervirulent M89 GAS bacteria

Having established that pRha-OMV antibodies raised in C57BL/6J mouse models are efficient in opsonising GAS pathogens, we investigated the effect of pRha-OMV immunised rabbit antibodies on the binding and killing of GAS strains. Increased rabbit pRha-OMV IgG antibody deposition was observed on newly emerged *S. dysgalactiae* subsp. *equisimilis* (SDSE) isolates, which have replaced their Group G Carbohydrate gene cluster with a functional Group A Carbohydrate cluster (SDSE_*gac*) [27]. The anti-pRha-OMV sera recognised and binds to SDSE_*gac* isolates Stg485 and Stg652 [Fig.5B]. Microscopic examination of the M89 strain treated with pRha-OMV rabbit serum displayed uniform cocci (long chains) revealing the classical structure of GAS [Fig.5C]. Furthermore, rabbit pRha-OMV antibody, but not rabbit pre-immune serum, was able to kill the hypervirulent strain M89 [Fig.5D]. Overall, these data confirm the ability of recombinantly produced pRha to induce type-specific functional antibodies that aid in promoting the recognition and killing of the M89 hypervirulent strain.

## Discussion and concluding remarks

Several recent studies have addressed the production of universal GAS vaccine candidates targeting the GAS polyrhamnose backbone of the group A carbohydrate. These approaches focused on chemical extraction of natively produced pRha from GAS cells or chemical conjugation to a carrier protein [12, 42]. We explored an alternative production route, and, for the first time, we have succeeded in producing recombinant pRha embedded into *E. coli* OMVs and evaluated them as vaccine candidates against GAS. The data demonstrate that pRha-OMVs were effectively presented to the mouse immune system and triggered a strong immune response specific for the pRha carbohydrate when supplemented with an adjuvant. This effect agrees with the findings of a study investigating *E. coli* OMV decorated with β-(1→6)–linked poly-N-acetyl-D-glucosamine (PNAG) [43]. PNAG is a conserved polysaccharide antigenic component found in bacteria, fungi and protozoan cells. Immunisation studies on PNAG-OMV provoked polysaccharide specific IgG antibodies.

The prevalence of IgG subtypes has been widely studied within the GAS infection models. One of the most abundant subclasses of human IgG found against the GAS M proteins is IgG3 [44]. Notably, antibody isotype class switching from IgM to mouse IgG3 constant regions was proven to be driven by the frequent exposure to polysaccharide antigens [45, 46]. A strong IgG1 response was observed in humans exposed to polysaccharides from pneumococcal pathogens [35] and increased IgG2a and IgG2b levels were detected in mice exposed to M protein-based vaccines from *Streptococcus pyogenes* [47].

Previous studies have shown that immunisation of rabbits with the pRha backbone extracted from native GAS promotes opsonophagocytic killing of multiple GAS serotypes and also protects the animals against systemic GAS challenges [11]. Here, we observed that anti-pRha-OMV IgG promoted bactericidal killing of the hypervirulent M89 strain mediated by baby rabbit complement. Strikingly, recognition of SDSE_*gac* isolates by pRha-OMV antibodies broadened the potential protection of a pRha-based vaccine to target newly emerging SDSE isolates.

A major finding of this study is the increased expression of IL-17a stimulated by the pRha-OMV antigen in mouse splenocytes. Carbohydrates are known to trigger a T cell independent immune response and hence conjugating them with a carrier protein elicits T cell dependent immune response. Recognition of polysaccharides by CD4^+^ T cells were shown to induce activation and differentiation which can be analysed by measuring the IL-17A cytokine [48]. IL-17-mediated protective immunity via neutrophil recruitment has been shown against GAS [39], *Klebsiella pneumoniae* [49] and *Streptococcus pneumoniae* [50]. However, we cannot rule out the possibility of a role for endogenous *E. coli* antigens in inducing IL-17a, which needs further exploration.

In summary, we have demonstrated that the recombinantly produced pRha in *E. coli* OMVs induces robust IgG antibodies in mice and stimulates IgG binding to all tested GAS serotypes. Importantly, the antibodies efficiently kill the hypervirulent strain M89 implying possible IgG-mediated protection in *in vivo* models. The antigenic properties shared between GAS and *E. coli* cells requires further research, and possible avenues include producing detoxified LPS recombinant pRha-OMV and studying its immunogenicity in animal models. Future exploration of recombinantly-produced pRha-OMVs as a universal GAS vaccine candidate, as well as the conjugation of recombinant pRha to a carrier protein, remains to be investigated.

