## Supplementary material for "Recombinant Group A Carbohydrate backbone embedded into Outer Membrane Vesicles is a potent vaccine candidate targeting Group A Streptococcus from *Streptococcus pyogenes* and *Streptococcus dysgalactiae* subsp. *equisimilis*": Suppl Figure

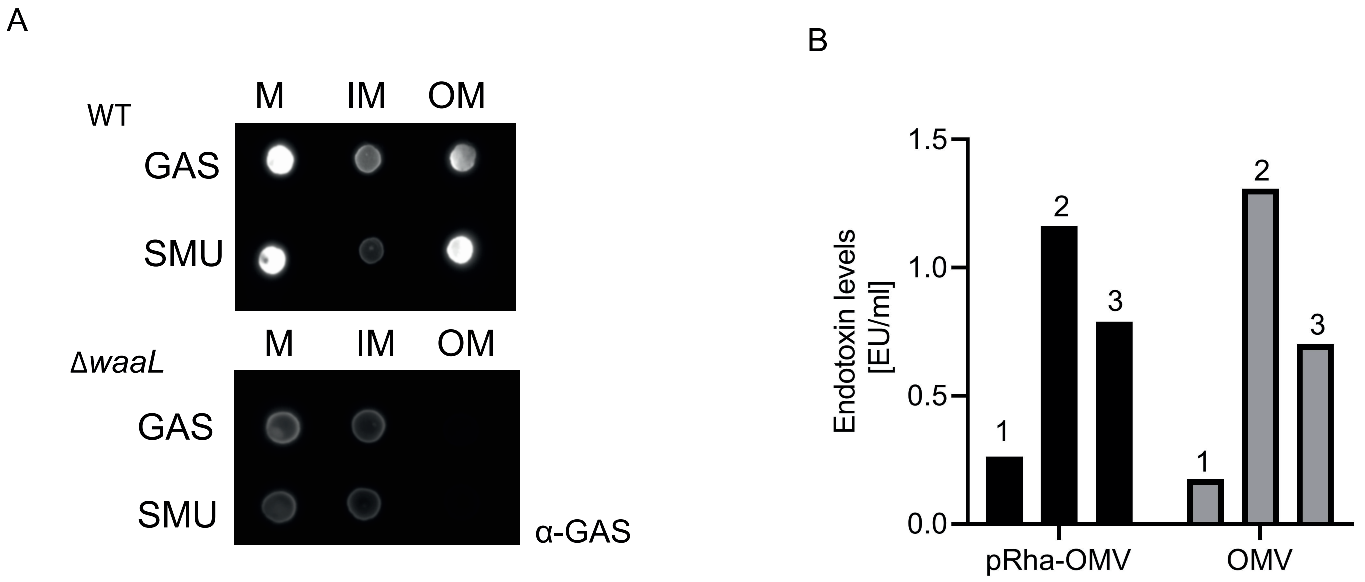


C

| Sample | Concentration /ml | Average Size |
|---|---|---|
| pRha-OMV | 4.0E+9 | 131.3 nm |
| OMV | 2.97E+7 | 153.3 nm |

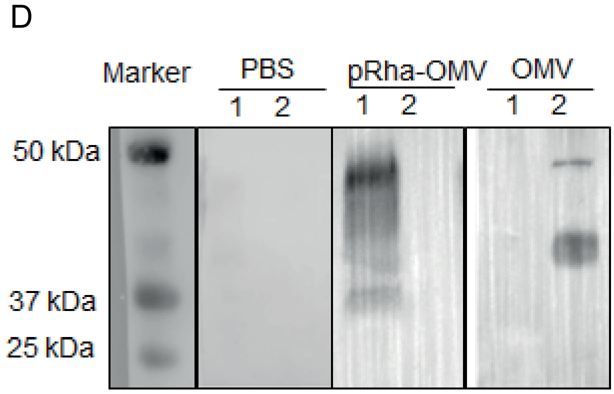


**Supplementary Figure 1: Supplemental data on the endotoxin levels, OMV particle size, pRha-OMV antibodies against *E. coli* producing polyrhamnose tested using Western blot analysis** A) Spot-Blot of the crude membrane (M), inner membrane (IM), and outer membrane (OM) of wildtype (WT) or ΔwaaL *E.coli* cells expresing the GAS and SMU gene clusters. B) Quantification of endotoxin levels in the purified pRha-OMV and OMV alone in first three doses used to immunise the animals. Number indicates the immunisations doses C) OMV concentration and the particle size in diameter (nm) from the purified *E. coli* producing pRha or *E. coli* OMV on their own were estimated using ZetaView nanoparticle tracking analyser with 670 nm laser. Samples were diluted in 1:4 using PBS. PBS was used as a control and subtracted from all samples. Data shown are mean ± SD of technical replicates D) Immunoblot analysis of whole *E. coli* OMV pRha lysate (1) or *E. coli* OMV alone (2) were blotted using pooled antiserum from the pRha-OMV and OMV or PBS vaccinated groups. Bound antibodies were probed using a secondary antibody of 1:2500 dilution of anti-rabbit IgG HRP and viewed using image lab software. Molecular mass markers are given in kilodaltons.
